# Frequent co-regulation of splicing and polyadenylation by RNA-binding proteins inferred with MAPP

**DOI:** 10.1101/2022.01.09.475576

**Authors:** Maciej Bak, Erik van Nimwegen, Ralf Schmidt, Mihaela Zavolan, Andreas J. Gruber

**Affiliations:** Computational and Systems Biology, Biozentrum, University of Basel, Klingelbergstrasse 50-70, 4056 Basel, Switzerland; University of Konstanz, Universitaetsstrasse 10, D-78464 Konstanz, Germany

## Abstract

Maturation of eukaryotic pre-mRNAs via splicing, 3’ end cleavage and polyadenylation is modulated across cell types and conditions by a variety of RNA-binding proteins (RBPs). Although over 1’500 proteins are associated with RNAs in human cells, their binding motifs, targets and functions still remain to be elucidated, especially in the complex environment of human tissues and in the context of diseases. To overcome the lack of methods for systematic and automated detection of sequence motif-guided changes in pre-mRNA processing based on RNA sequencing (RNA-seq) data we have developed MAPP (Motif Activity on Pre-mRNA Processing). We demonstrate MAPP’s functionality by applying it to RNA-seq data from 284 RBP knock-down experiments in the ENCODE project, from which MAPP not only infers position-dependent impact profiles of known regulators, but also reveals RBPs that modulate both the inclusion of cassette exons and the poly(A) site choice. Among these, the Polypyrimidine Tract Binding Protein 1 (PTBP1) has a similar activity in glioblastoma samples. This highlights the ability of MAPP to unveil global regulators of mRNA processing under physiological and pathological conditions.

## Introduction

Splicing and 3’ end processing of nascent RNAs are crucial steps in eukaryotic pre-mRNA processing, also responsible for transcriptome diversification through the generation of isoforms. Both processes are modulated by various RNA-binding proteins (RBPs), whose expression varies across tissues. To date, a few dozen regulators have been described to modulate splicing and/or 3’ end processing ^1,2^. However, the functions of the vast majority of the more than 1’500 RBPs encoded in the human genome remain to be unraveled ^3^. Recent work has started to identify RBPs that act at multiple stages of pre-RNA processing ^4–6^, but a tool for the systematic discovery of such regulators is lacking. To address this issue, we have developed MAPP (Motif Activity on Pre-mRNA Processing), a computational method that enables the identification of RBP-specific sequence motifs that can explain global patterns of pre-mRNA processing observed in RNA-seq data. MAPP makes use of a powerful functional concept that we have previously exploited in our KAPAC tool ^7^, namely explaining relative expression levels of transcript isoforms across samples with sequence motifs located throughout nascent transcripts. In contrast to KAPAC, MAPP provides an end-to-end solution to the inference of motifs, known or not to bind specific RBPs, that influence splicing, 3’ end cleavage or both processes.

We benchmarked MAPP using data sets in which RBPs with well-characterized impact on splicing and/or 3’ end processing have been overexpressed or depleted by siRNA-mediated knock-down, showing that MAPP identifies not only the correct sequence motif, but also the binding site position-dependent impact of the RBP on RNA processing, at a fine level of detail. We confirmed a repressive function of heterogeneous ribonucleoprotein C (HNRNPC) on both the inclusion of cassette exons and the processing of poly(A) sites, effects shared by another pyrimidine-rich element-binding protein, Polypyrimidine Tract Binding Protein 1 (PTBP1). The latter also explains best the global dysregulation of pre-mRNA processing ^8^ that occurs in glioblastoma cancers, not only at the level of polyadenylation, as we reported before ^7^, but also at the level of splicing. We also highlight the position-specific effect of RNA Binding Fox-1 Homolog 1 (RBFOX1) on splicing and that its effect depends on the location of the binding sites within the nascent transcript. After benchmarking MAPP on these three RBPs with known impact on pre-mRNA processing, we turned to the >400 RBP knockdown experiments publicly available from the ENCODE project, aiming to systematically determine RBPs with position-dependent regulatory impact on pre-mRNA processing and infer their sequence specificity. This approach identified multiple RBPs whose specific impact on pre-mRNA processing has not been reported to date, among which various pyrimidine motif-binding proteins seemingly explain changes in both exon inclusion and the poly(A) site choice. In summary, in this study we inferred the RNA processing impact map of a large number of RBPs and identified those that regulate both splicing and 3’ end processing. Also, we deliver a novel computational approach that enables a comprehensive characterization of the regulatory motifs and their position-specific impact on splicing and/or 3’ end processing within cellular systems of interest. “*MAPPing*” RNA-seq experiments will thus enable the identification of regulators that can explain global changes in pre-mRNA processing observed in health and diseases.

## Results

### A model for inferring the positional impact of splicing and 3’ end processing regulators

Whereas RBPs have long been known to orchestrate pre-mRNA splicing (e.g. ^6^), their impact on 3’ end processing has only recently started to become apparent ^4,9–11^. In particular, the question of whether RBP regulators act on splicing and 3’ end processing in a coordinated manner has drawn much attention in recent years ^7,11^. A bottleneck in addressing this question is that compared to other types of regulators, such as transcription or epigenetic factors, the fraction of RBPs with well-characterized binding specificities is relatively minor. In addition, even for those RBPs for which binding specificities have recently been characterized by high-throughput sequencing, their impact and mode of action on pre-mRNA processing remain speculative. To address such questions, we have developed MAPP (Fig. 1).

**Fig. 1.**
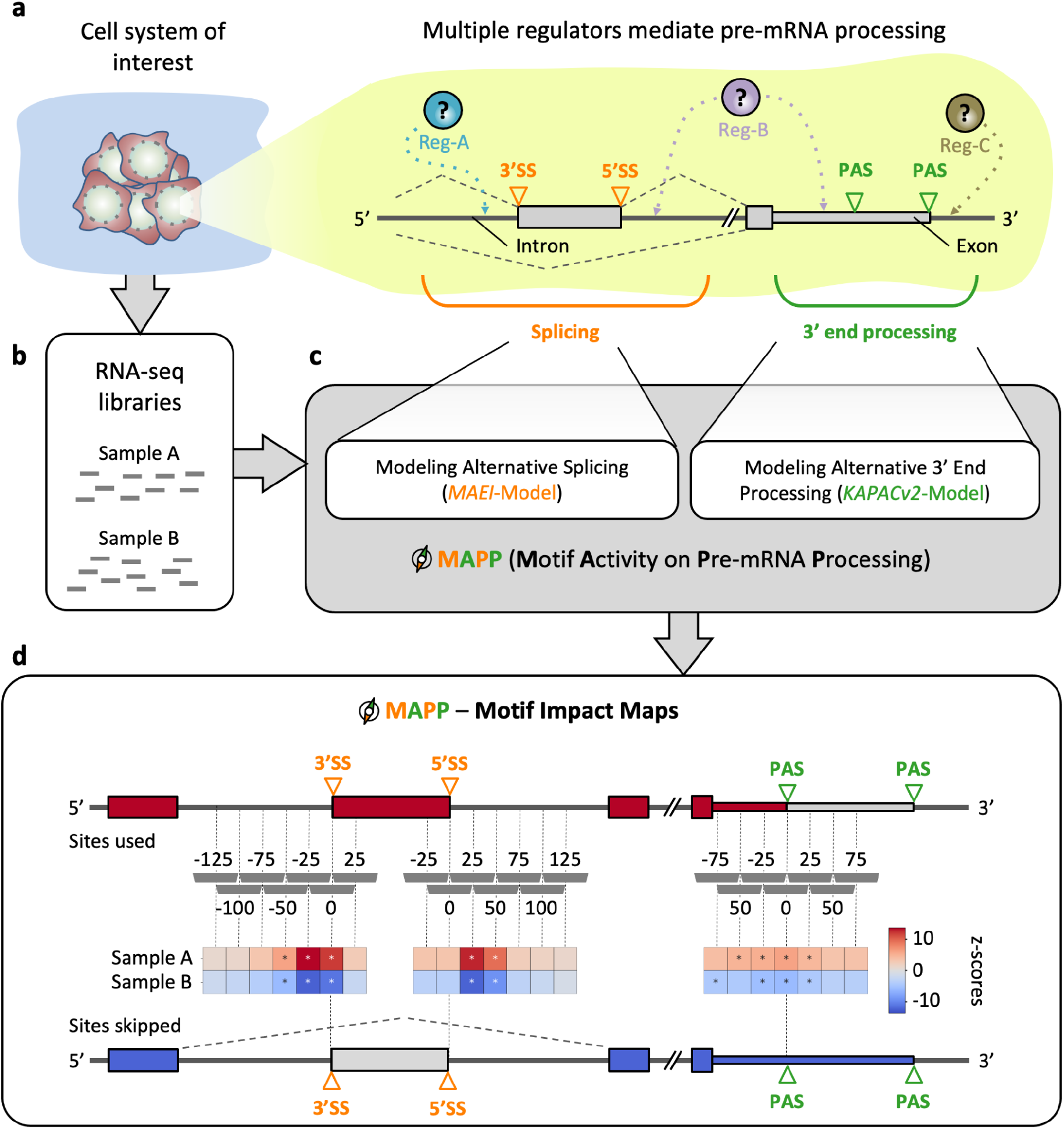
Inferring splicing and 3’ end processing motif impact maps using MAPP. **a |** Sketch illustrating how regulators (Reg) bind pre-mRNAs to influence the usage of splice sites (SS) and / or poly(A) sites (PAS). **b |** RNA-sequencing (RNA-seq) libraries are available or can be created for most cellular systems of interest. **c |** MAPP is an automated tool for analyzing the splicing and 3’ end processing patterns inferred from RNA-seq data with the MAEI (Motif Activity on Exon Inclusion) and KAPACv2 (K-mer Activity on PolyAdenylation site Choice version 2) models, respectively. **d |** MAPP infers regulatory motifs for RBPs and reports detailed maps of their position-dependent impact on cassette exon inclusion and poly(A) site usage, respectively.

MAPP includes a component designed to infer the position-dependent activity of sequence elements on splice site (SS) choice (see Methods: MAEI), and a component that infers the activity of motifs on poly(A) site choice, the latter building on our previously-described KAPAC approach ^7^. While similarly to KAPAC, MAPP considers the entire space of sequence motifs, modeled as k-mers, that could impact pre-mRNA processing, its functionality is more general, as it can also work with position-dependent weight matrices (PWMs, see Methods: KAPACv2) representing known binding specificities of RBPs. The two components model changes in exon inclusion and poly(A) site usage across genes as functions of the motif counts within regions located at various distances relative to these events (see Methods). More specifically, given RNA sequencing data from a cellular system of interest (Fig. 1a,b), MAPP first infers the level of inclusion of alternatively spliced exons and the usage of distinct poly(A) sites (Supplementary Figures S1 and S2). Then, the MAEI and KAPACv2 models are fitted to the corresponding event data to identify sequence motifs that can explain global splicing and 3’ end processing patterns, respectively (Fig. 1c). By applying the models to sequence windows (for all our analyses unless specified otherwise: 50nt in length, slided by 25nt) located at specific distances relative to pre-mRNA processing sites, position-dependent activity z-scores are inferred for each motif. MAPP ranks the sequence elements based on their significance (Fig. 1d) and reports the position-dependent z-scores in the form of impact maps ^7^. Thus, MAPP infers the impact maps both for motifs known to correspond to specific regulators, as well as for motifs that have not yet been linked to a specific RBP.

### MAPP recovers the position-dependent impact of known regulators

To validate MAPP, we applied it to data sets from experiments where proteins with a known effect on splicing/polyadenylation were perturbed. We started with the known HNRNPC regulator and found that the sequence motif most significantly associated with both the measured changes in exon inclusion as well as poly(A) site usage is penta-U, the motif known to be bound by HNRNPC ^12–14^ (Fig. 2a). The PWM representing this motif had the largest combined z-score out of the 344 PWMs that we curated from the ATtRACT database (see Methods). Furthermore, MAPP recovers the impact map expected for the HNRNPC RBP. That is, in control (CTRL) samples, where the expression of HNRNPC is high, MAPP infers a repressive effect (marked as blue squares) on 3’ splice site (3’SS), 5’ splice site (5’SS) and polyadenylation site (PAS) processing. Conversely, these sites are processed and thus the activity of the penta-U motif is positive in the knock-down cells, where the expression of HNRNPC is low. These results are supported by a multitude of studies (e.g. refs ^4,14,15^). Additionally to the impact maps generated based on transcriptome-wide data we have selected 3’SS, 5’SS and poly(A) sites of the 200 exons whose inclusion/processing changes most (in either direction) or changes least upon HNRNPC knock-down and constructed position-dependent densities of predicted HNRNPC binding sites within the ranges we use to fit MAPP on. The resulting density profiles of the top 200 most changing mRNA processing targets are consistent with the impact maps, indicating that the changes in RNA maturation are due to interactions of HNRNPC with the RNAs.

**Fig. 2.**
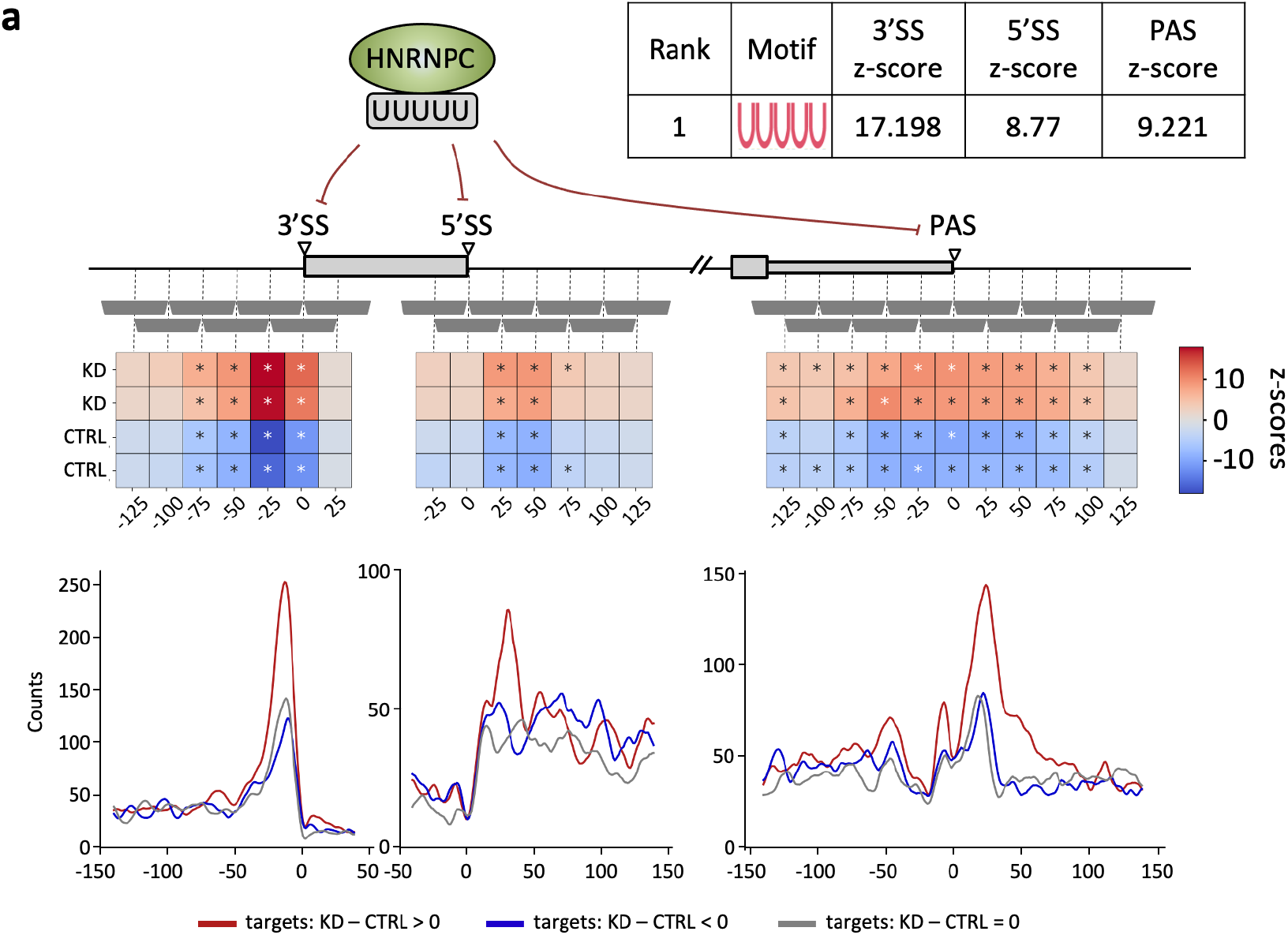
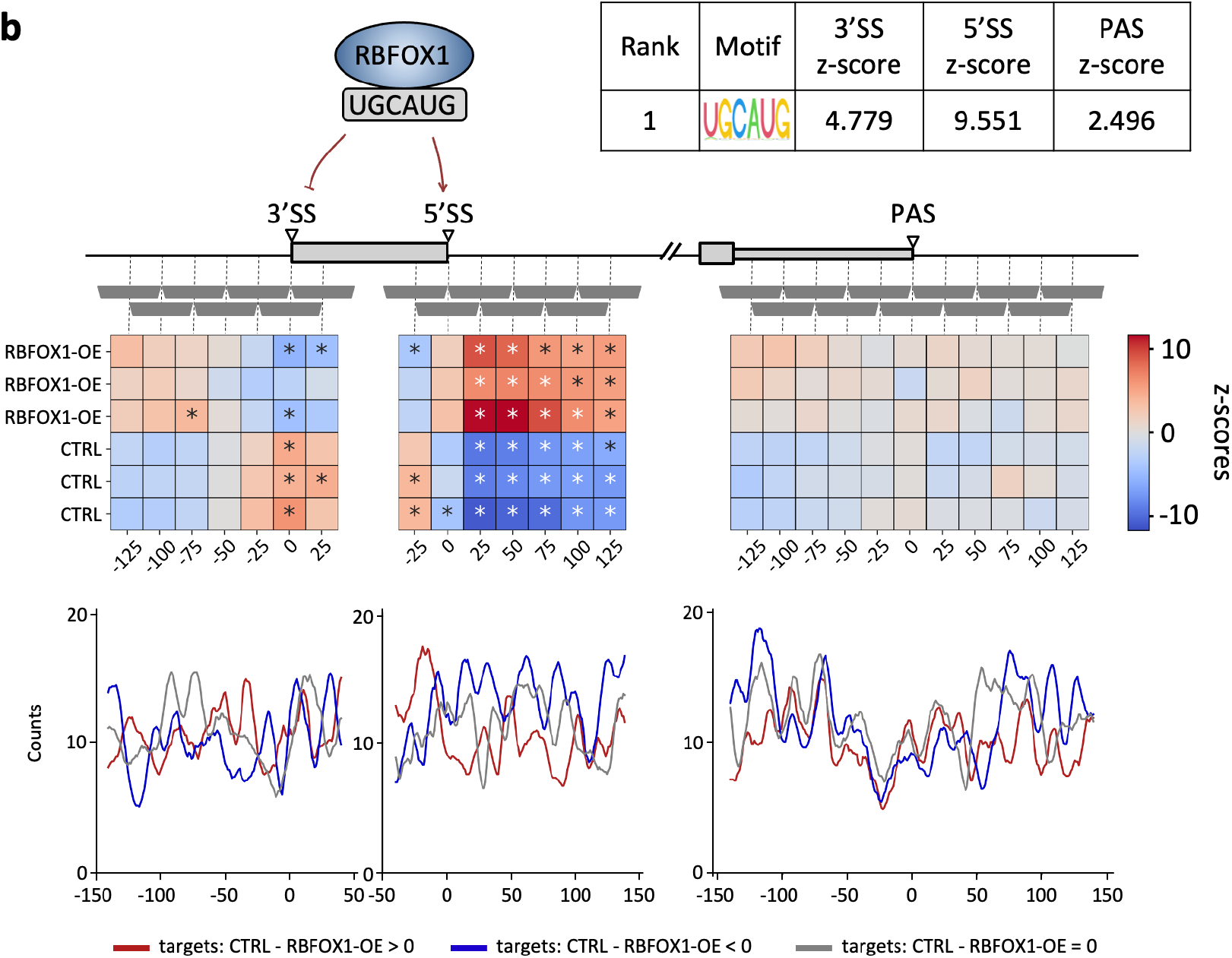
MAPP infers the known regulatory impact of the HNRNPC and the RBFOX1 RBPs on splicing and 3’ end processing from RBP expression perturbation data sets. **a |** Top panel: z-scores of activity changes in the vicinity of 3’ splice sites (3’SS), 5’ splice sites (5’SS) and poly(A) sites (PAS) inferred from a HNRNPC knock-down data set. The PWM with the highest inferred combined z-score of all PWMs available in ATtRACT database has the penta-U motif as consensus. By fitting the splicing and 3’ end processing models of MAPP to overlapping windows (horizontal gray bars) located at specific distances relative to splice and poly(A) sites, position-dependent activity z-scores are inferred. Statistically significant z-scores are marked with an asterisk. Bottom panel: Smoothened (+/− 5 nt) densities of the inferred HNRNPC binding motif in the vicinity of the top 200 3’SS, 5’SS and PAS, whose usage is most upregulated (red), most down-regulated (blue) or does change least (gray) upon HNRNPC knock-down. **b |** Top panel: MAPP results as described in **a**, but here applied to a RBFOX2-deficient HEK293 cell line with induced expression of RBFOX1 which is known to regulate splicing at 5’SS by binding to UGCAUG sequences ^17^. Bottom panel: Densities as in **a**, but for the RBFOX1 binding motif in the vicinity of the top 200 3’SS, 5’SS and PAS, whose usage is most upregulated (blue), most down-regulated (red) or does change least (gray) upon RBFOX1 expression.

We next turned to a well characterized splicing regulator, the RBFOX1 RBP. Analyzing data from an experiment where the RBFOX1-dependency of exons was determined in RBFOX2-deficient HEK293 cells in which RBFOX1 was inducibly expressed from a Flp-in locus ^16^, MAPP ranked the previously described RBFOX1-binding sequence, UGCAUG, as the most significant in explaining exon inclusion and inferred its activating activity when located downstream of 5’SS ^17^. MAPP also highlights the opposite activity near the 3’SS, where RBFOX1-binding motifs are associated with reduced exon inclusion. While this repressive effect appears to be weaker compared to the activating effect of motifs located downstream of 5’SS, it is interesting to infer simply from the RNA-seq data that RBFOX1, like other RBPs ^18^, can have opposing impacts depending on the location of a binding site. These results show that MAPP enables fine grained insights into the binding-specificity and position-dependent impact of RBPs on splicing and 3’ end processing.

### The PTBP1 RBP binding motif explains splicing and 3’ end processing changes in glioblastoma

As key factors in the post-transcriptional regulation of gene expression, RBPs have been reported to play an important role in numerous diseases, including cancer ^19^. Moreover, various groups have reported that the 3’ UTRs of transcripts become shortened upon tumorigenesis ^20–22^, thus presumably escaping the RBP-mediated regulation. In a previous study we have uncovered that PTBP1 best explains the global remodelling of 3’ UTR length in glioblastoma ^7^. Thus, to further validate MAPP, we applied it to PTBP1/2 double knock-down data. As expected, MAPP inferred that PTBP1 has a repressive activity on the processing of 3’SS and to some extent 5’SS, as PTBP1-binding motifs located in these regions are associated with reduced exon inclusion when the level of PTBP1 is high, i.e. in control conditions (Fig. 3a). Furthermore, PTBP1 also has a repressive effect on polyadenylation, as we have reported before ^7^. Turning to glioblastoma cancer data from The Cancer Genome Atlas (TCGA) ^23^, we recovered the PTBP1-binding motif as having the highest activity on polyadenylation (Fig. 3b). Furthermore, MAPP infers that changes in exon inclusion are also best explained by the PTBP1-binding motif, with a position-dependent activity that matches the profile obtained from the PTBP1/2 knock-down data. These results strengthen the case for PTBP1 as a main regulator of pre-mRNA processing in glioblastoma, a cancer where PTBP1 is highly overexpressed (Fig. 3b).

**Fig. 3.**
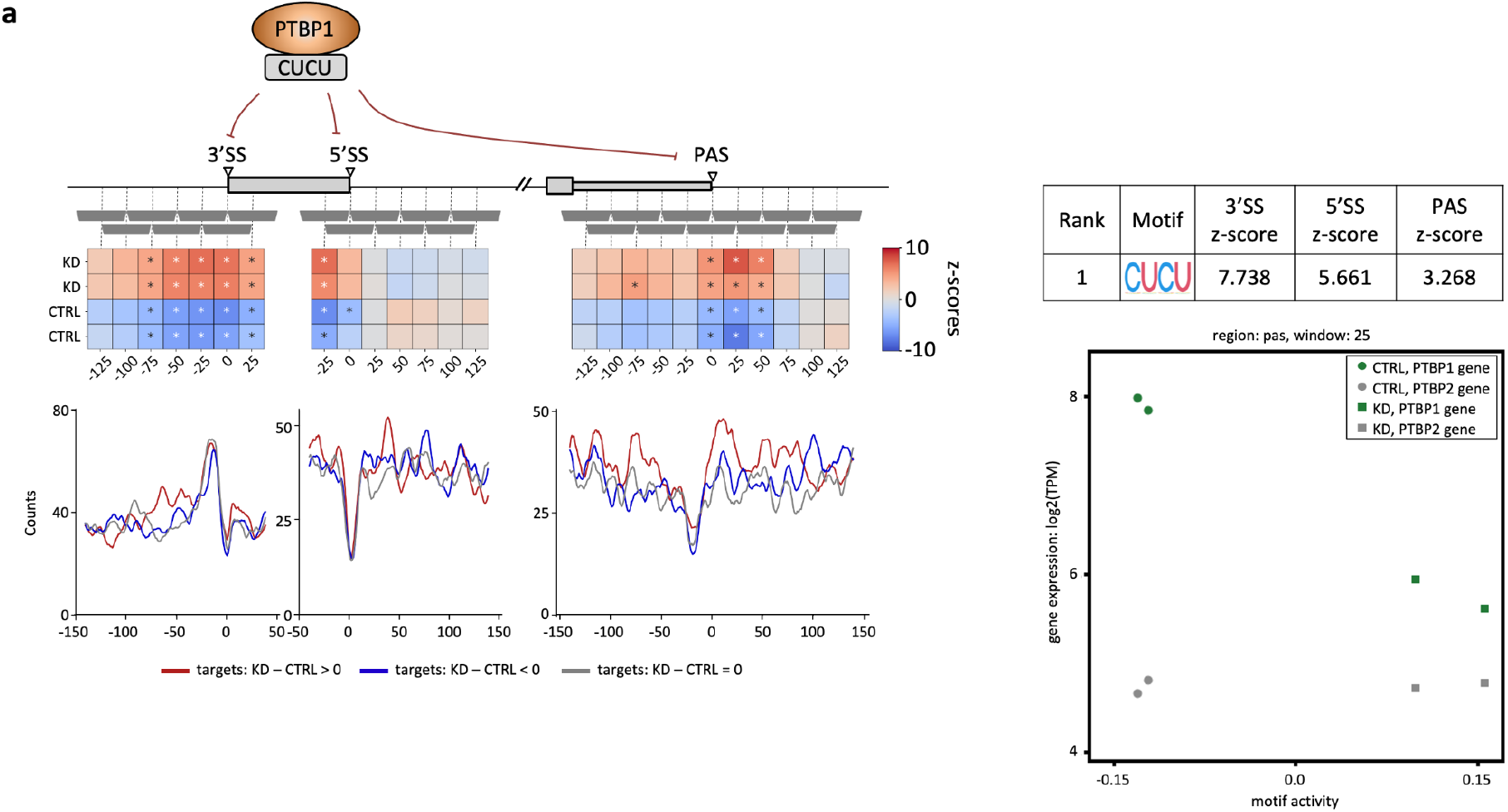
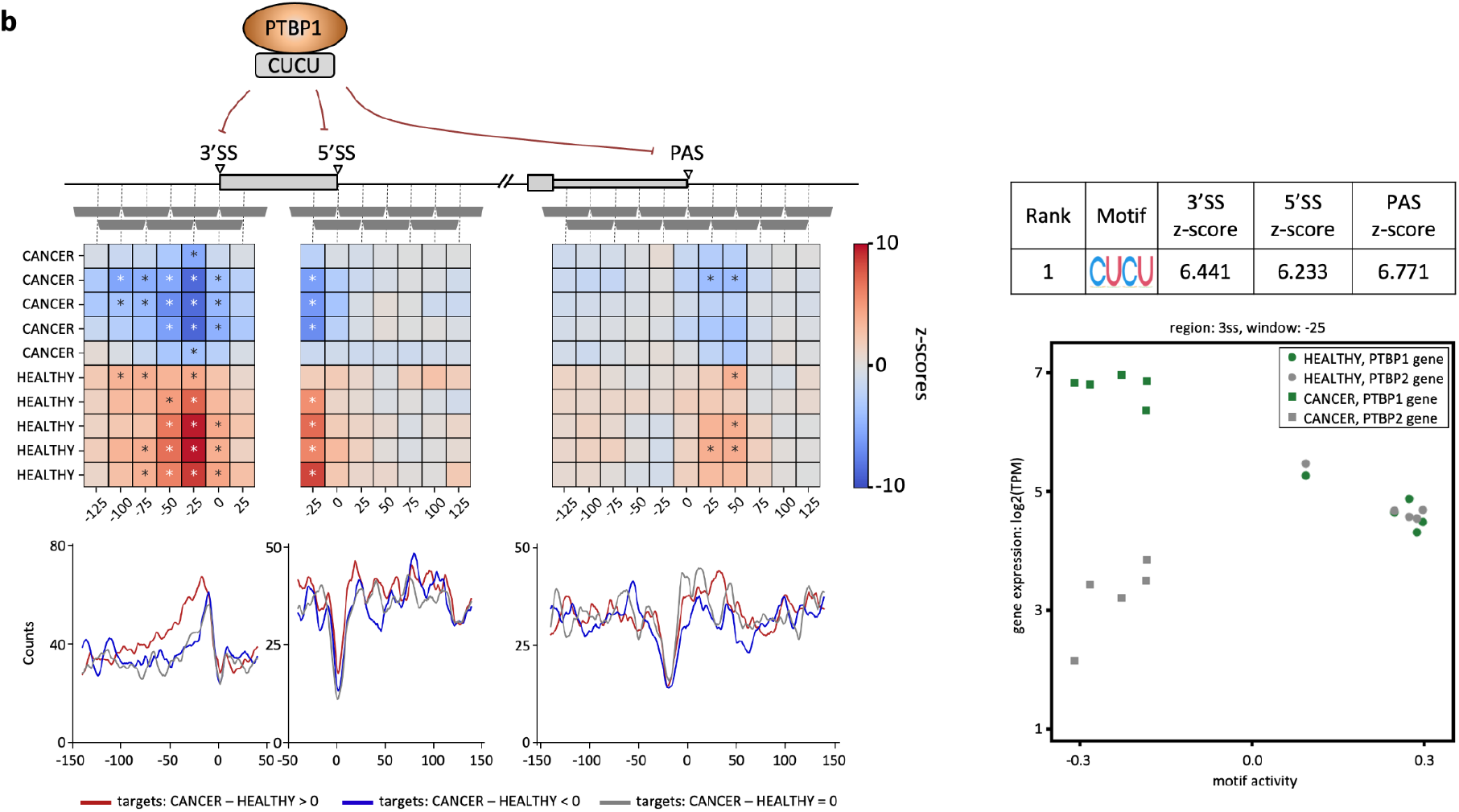
Splicing and 3’ end processing activities of the PTBP1 RBP in. **a |** cells depleted of both PTBP1 and PTBP2 by siRNA-mediated knockdown, and **b** | glioblastoma samples. Panel structure is the same as in Fig. 2, and in addition, scatter plots of the activities inferred for the top ranked motif (shown in the tables) versus mRNA expression levels of PTBP1 and PTBP2 (log2 TPM) across samples are shown.

### MAPP infers the binding specificities and detailed impact maps for pre-mRNA processing regulators in the ENCODE data sets

Next, we used the large array of RBP knock-down datasets available from the ENCODE project to comprehensively infer the sequence specificity, position-dependent impact and activating or repressive mode of action of human RBPs on pre-mRNA processing. Applying MAPP to 456 RBP knock-down experiments available in ENCODE we found that the tool is indeed able to identify the motif known from the ATtRACT database to correspond to the protein whose expression was altered in the experiment. Fig. 4 shows summary results for samples (see Methods) for which the RBP corresponding ATtRACT-provided PWM was ranked among the top 5 most significant motifs. Interestingly, RBP motifs generally have the same type of activity on exon inclusion - activating or repressive - when located at both 5’ and 3’ splice sites, the only exception being the QKI motif, inferred to repress processing when located around the 3’SS and promote it when located around 5’SS. Notably, the general splicing factor SRSF1 is inferred to promote splice site processing, as expected, while many RBPs that act as splicing modulators (e.g. HNRNPC, PTBP1, HNRNPK) are inferred to have a repressive role. A notable exception is the Poly(rC) Binding Protein 1 (PCBP1) RBP, also estimated to promote the splicing of target exons.

**Fig. 4.**
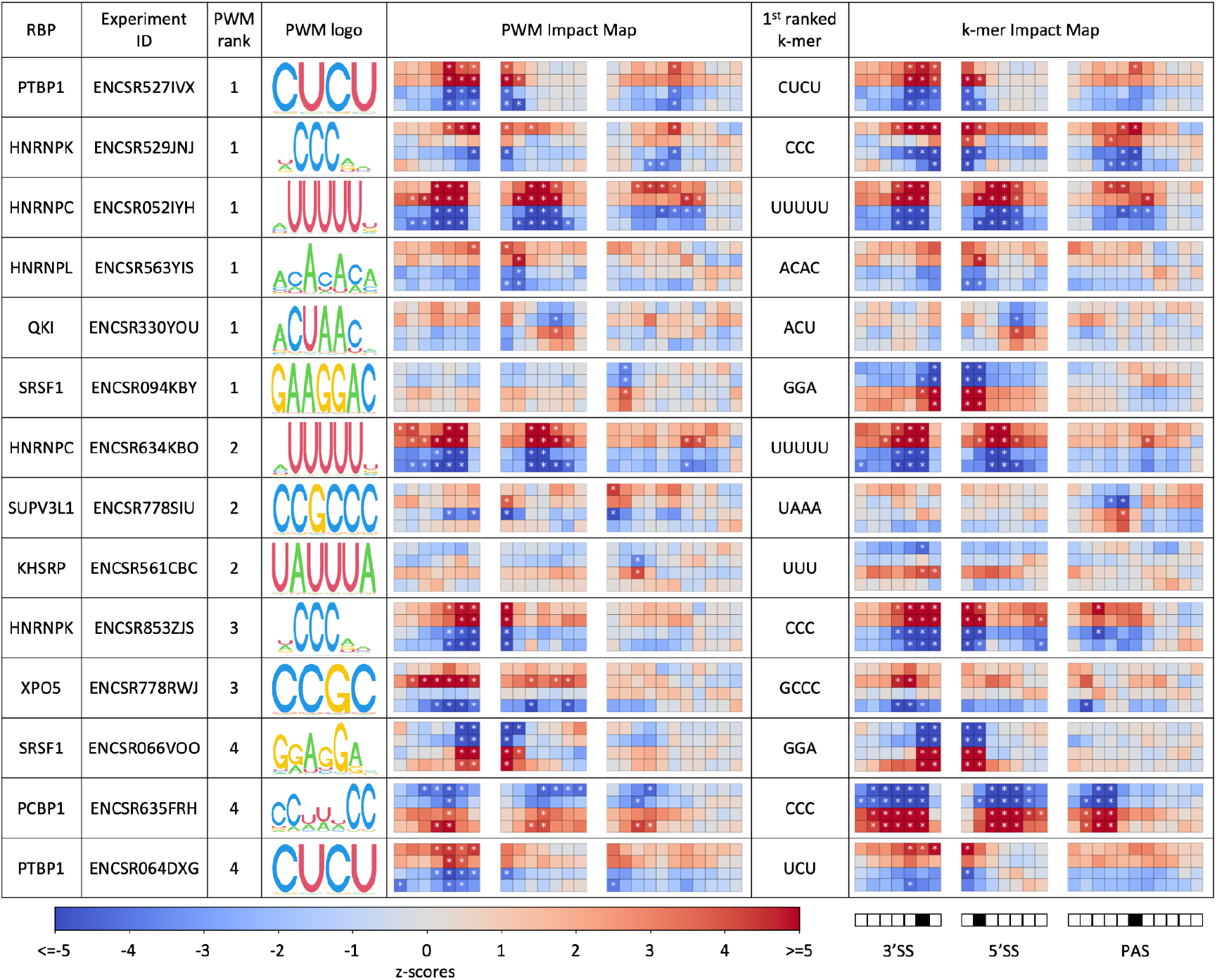
MAPP reveals the concerted impact of pre-mRNA processing regulators on splicing and polyadenylation. For each RNA-binding protein we determined the first motif (in the order of MAPP-provided significance) that is assigned in ATtRACT to the RBP that was depleted in each experiment. Column 3 shows the rank of that motif given by MAPP. The table contains RBPs where the known binding motif was among the top 5 reported by MAPP. The activity profiles are shown similarly to those in Figs. 2 and 3, the top two rows indicating the knock-down and the bottom two the control conditions. Windows within regions around 3’SS, 5’SS and PAS are set to the same ranges as for our previous analyses (Figs. 2 and 3). The central window sliding through a given RNA processing site (−25nt,+25nt) was marked as black square in the legend. Furthermore, in addition to the PWM-based MAPP runs, we have carried out a similar analysis exploring all possible k-mers of length 3 to 5. The top-ranked k-mer is reported for each experiment alongside the corresponding PWM result.

As the AtTRACT-provided PWMs corresponding to the perturbed proteins were not always the most significant motifs in explaining the RNA processing alterations, we also ran MAPP in the k-mer mode, to determine which sequence elements explained best the observed changes. Interestingly, for some data sets the results of the k-mer analysis were both more significant and more stable between replicate knock-down experiments. For instance, for the two independent knock-down experiments of the SRSF1 splicing factor, two replicates per condition each, MAPP inferred two distinct AtTRACT-reported SRSF1-associated PWMs, ranked as the first and fourth most significant PWM in the two experiments, respectively. In contrast, running MAPP in k-mer mode consistently inferred the core GGA motif to be most significant in both experiments. The inferred impact maps further confirm that the k-mer based MAPP results are highly consistent, whereas the PWM-based results can differ between experiments.

MAPP also reveals further regulatory patterns (Fig. 4), namely that half of the RBPs shown in the table impact both splicing and polyadenylation. Moreover, the RBPs that impact both processes act consistently as either activators or repressors of both exon inclusion and poly(A) site usage. This may hint to a concerted regulation of alternative terminal exons by individual regulators, but it must go beyond the regulation of terminal exons, because the motif has similar activity around the 5’SS, which does not occur in the terminal exons. Finally, MAPP also indicates that RBPs with similar sequence specificity can exert their regulatory roles by binding to the pre-mRNA in different positions relative to the pre-mRNA processing sites. For instance, both the PCBP1 and the HNRNPK RBPs bind a ‘CCC’ sequence element to regulate splicing and polyadenylation. However, while the impact of HNRNPK seems to be focused on the immediate vicinity of processing sites, PCBP1 appears to activate splicing from a broader region.

While the proteins with known PWMs shown in Fig. 4 have been implicated in splicing before, we were especially interested in cases where MAPP identified a clear k-mer, but not a PWM of a known regulator as being able to explain the pre-mRNA processing changes. The most compelling example appears to be the PUF60 RBP. ENCODE provides knock-down experiments for this protein in two cell lines: K5643 (*ENCSR558XNA*) and HepG2 (*ENCSR648BSC*). The two experiments that exhibit the most significant MAPP results consistently infer a highly similar U-rich sequence element (Supplementary Fig. S3), which is also the motif inferred to be bound by PUF60 *in vitro*, in RNA Bind’n-seq experiments (encodeproject.org). MAPP also charts the impact of PUF60 on the 3’ splice site at a fine level of detail, which is consistent with a previous report of PUF60 activating exon inclusion by binding to U-rich regions upstream of 3’ splice sites ^24^. MAPP pinpoints the activating impact of PUF60 to the intronic region of ~25-0 nts upstream of the 3’ splice site, which is much more position-specific than other regulators mentioned above. These results illustrate the utility of MAPP in elucidating the regulation of pre-mRNA processing by RBPs.

## Discussion

By binding to sequence elements in transcripts, RNA-binding proteins regulate gene expression at co-transcriptional and post-transcriptional levels. In particular, they can affect both splicing and 3’ end processing, key steps in the maturation of mRNAs. Additionally, the interaction of RBPs with mature mRNAs can regulate the transport, localization and translation of these mRNAs ^25^. Understanding of the global and concerted effect of various RBPs on the cellular transcriptome is undoubtedly key to understanding how gene expression is dysregulated in various pathological conditions, including cancer ^26,27^. In this study we presented a novel computational approach for inferring the regulatory impact of various RBPs on splicing and 3’ end processing.

We have validated our method on data pertaining to proteins with well-established roles in splicing and/or polyadenylation. Specifically, the MAPP-inferred activities of the HNRNPC RBP are in line with its previously reported role in preventing exonization of cryptic *Alu* elements ^15,28^. Many *Alu* elements have evolved to become cassette exons, and the potentially deleterious inclusion of these exons needs to be tightly regulated. The impact maps constructed by MAPP are fully consistent with this role of HNRNPC (Fig. 2a). MAPP also recovers the previously-noted position-specific regulation of exon inclusion by RBFOX1 (Fig. 2b), whereby binding of RBFOX1 upstream of cassette exons results in their exclusion, while binding downstream of such exons promotes their inclusion ^29^. While our results are consistent with this model, they also provide higher granularity in the binding site position-dependent effects of RBFOX1. Specifically, they indicate a higher impact of the downstream, inclusion-promoting sites. Furthermore, MAPP indicates that binding sites that are located further upstream in the introns also have an overall inclusion-promoting effect, consistent with an earlier report ^17^. Thus MAPP provides direct and broad insight into the activity of RBP binding sites from individual RNA-seq data sets, without a need for stratifying the data or determining the binding sites with methods such as crosslinking and immunoprecipitation. MAPP’s position-dependent activity maps thus enable an efficient and improved understanding of how RBPs exert their roles both globally and on individual targets.

MAPP analysis of glioblastoma samples yields impact maps that strikingly resemble those obtained from PTBP1/2 knock-down data (Fig. 3a,b). That the PTBP1-binding motif is also the RBP-binding motif that best explains the splicing deregulation observed in these samples strengthens our previous hypothesis that PTBP1 is a key regulator of RNA processing in this cancer ^7^. Identification of RBPs that broadly impact mRNA processing in specific conditions and in particular in individual cancers is highly relevant, as it can provide novel entry points for the development of therapies. Targeting of mRNAs and mRNA-RBP interactions with antisense oligonucleotides ^30,31^ or small molecules ^32^ holds much promise for medical applications. As MAPP is a fully automated workflow, the task of identifying regulators of pre-mRNA processing from novel RNA-seq datasets is considerably facilitated.

Our analysis of RBP knock-down data sets available from the ENCODE project revealed that multiple regulators affect exon inclusion and/or the poly(A) site usage (Fig. 4). Once again, thanks to the sliding window approach of MAPP we found that distinct regulators differ not only in their role (which for most of the individual RBPs seem to be the same in the two processes, i.e. to either enhance or repress) but also in their position specificity. For example, MAPP not only highlighted the opposite effect of two proteins, HNRNPK and PCBP1, which bind the same ‘CCC’ sequence, on cassette exon inclusion, but also that the distance range of their impact differs: PCBP1 acts more broadly in the introns flanking the cassette exon, while HNRNPK acts in a more focus manner, at the exon-intron boundaries. Of course, a knock-down is an artificial condition, while within tissues multiple RBPs likely vary in concentration in a concerted manner, leading to more complex patterns of regulation of mRNA maturation ^33^. Nevertheless, understanding the impact of individual RBPs based on individual knock-down experiments will facilitate the interpretation of their effect in tissue context.

Other groups have investigated binding site location-dependent effects of RBPs, specifically proposing the concept of “RNA maps” ^34^, which summarize the density of RBP binding sites in the vicinity of various types of landmarks (exon and transcript boundaries), where RBP exert regulatory roles. For instance, binding of the Nova RBP upstream of a cassette exon is associated with the skipping of that exon, while the binding downstream of the cassette exon is associated with the exon inclusion. The impact maps that MAPP constructs provide complementary information. They do not rely on direct information about the location of the binding site of the RBP (usually obtained with CLIP), nor on specific thresholds for defining regulated events such as exon inclusions. Rather, MAPP makes use of the quantitative information in the inclusion level of each exon or PAS as well as in the number of predicted binding sites in the vicinity of these exons. As a result, MAPP provides quantitative information about the impact of motifs on RNA processing, circumventing issues regarding the coverage of the binding sites by CLIP in targets with different levels of expression. Also interesting to note is the increasing use of massively parallel assays for exploring the dependence of RNA processing on specific motifs ^35^. These provide information more analogous to MAPP’s impact maps, but are limited to a small number of conditions, a small number of targets and regions within these targets, and have been so far used to characterize general principles of RNA processing. In contrast, MAPP’s utility comes primarily in exploring a broad range of conditions and identifying condition/tissue-specific regulators. Finally, deep learning models have also started to be used to explain patterns of RNA processing ^36,37^. The aim of these approaches is generally to predict the occurrence of specific events (e.g. the level of exon inclusion) in different samples. This is different from MAPP’s aim of inferring regulators of these processes. Thus, MAPP extends the RNA biologist’s toolbox to enable the functional characterization of RBP-RNA interactions at an increasing level of detail.

In conclusion we developed a powerful computational approach to identify regulators of splicing and 3’ end processing, which are frequently coordinated. MAPP has been developed using modern principles of software engineering, facilitating further development by a broad community of developers.

## Methods

### Datasets

We validated our method on publicly available RNA-seq data with perturbed levels of RBPs with known impact on splicing, and, to some extent, polyadenylation. The full list of samples’ and records’ IDs is included as Supplementary Table S1. Similarly, Supplementary Table S2 lists all RNA-Seq data sets related to RBP knock-downs we have obtained from the ENCODE project. To apply MAPP we require that samples meet minimal criteria of quality. For example, we require a sufficiently high Transcript Integrity Number (>50, typically > 70) ^38^, high proportion of uniquely mapped reads (>0.95), high proportion of high-quality mapped reads (>0.85; following RNA-SeQC’s documentation: proportion of properly paired reads with less than 6 mismatched bases and a perfect mapping quality out of all mapped reads), low level of rRNA contamination (<0.05) and low proportion of reads mapped to intergenic regions (<0.1), as reported by RNA-SeQC ^39^. BAM files with mapped RNA-seq reads of normal and tumour sample pairs from TCGA were obtained from the Genomic Data Commons (GDC) data portal ^40^. The selection of normal-tumour pairs from glioblastoma data was done as described previously ^7^.

### MAPP

MAPP, standing for **M**otif **A**ctivity on **P**re-mRNA **P**rocessing, enables the identification of previously unknown RBP binding sites which can explain global patterns of pre-mRNA processing in a cellular system, including healthy and diseased states. The tool is implemented as a modular snakemake workflow ^41^ with distinct standalone sub-workflows dedicated to separate functionalities. These are: RNA-Seq data preprocessing, selection of cassette exons, selection of tandem poly(A) sites, quantification of exon inclusion, quantification of poly(A) site usage, generation of matrices motif counts (PWMs/k-mers) in each window around each site, MAEI model (splicing), KAPACv2 model (polyadenylation), and summary of results. Each of these modules is described in more detail in Supplementary Materials (see Supplementary Methods). We support two distinct software technologies in order to ensure reproducible research: Conda environments ^42^ and Singularity containers ^43^.

#### MAEI

MAEI, which stands for **M**otif **A**ctivity on **E**xon **I**nclusion. MAEI is a novel model designed to infer the positional impact of various short sequence motifs on the differential inclusion of cassette exons. The input to the model is a quantification of transcript isoform expression along with the transcript annotation, and the matrix of motif counts in each window around signals of interest (5’SS, 3’SS). The motifs can either be specified as PWMs or k-mers. We model the inclusion level of every exon *e* in every sample *s* with a logistic function: *Θ _e,s_* = *e ^z^* / (1 + *e ^z^*), where *z* = *b _s_* + *c _e_* + *N _e,m_* * *A_m,s_* includes the model parameters: *b_s_* - the baseline inclusion of all exons in sample *s*, *c_e_* - the baseline inclusion of exon *e* in all samples and *A_m,s_* - the ‘activity’ of a motif *m* in a sample *s*. *N_e,m_* denotes the number of binding sites of a motif *m* in the proximity of exon *e*, which is either given by the sum of site probabilities predicted with the PWM or the raw k-mer counts). We further write down a likelihood to observe exon inclusion fractions in the whole data set as quantified and fit model parameters with an EM algorithm to obtain their maximum likelihood estimates. Per-sample motifs’ activities are then transformed to activity z-scores and presented visually on the impact maps. For motif ranking we implemented two separate strategies, suitable for distinct types of analyses: average z-score ranking (to identify regulators that act broadly across samples) and maximum z-score ranking (to identify regulators with strong impact but potentially in a small number of samples). Please see Supplementary Methods for more details on the calculations.

As a method of background correction we apply a Gaussian/Uniform mixture model to the per-sample distribution of activity z-scores and, again, find its maximum likelihood parameters by an EM method. We then standardize these z-scores by the inferred Gaussian parameters and transform them into p-values from a standard normal distribution. Statistical significance is later assessed upon Bonferroni-correction of such obtained p-values. Please see Supplementary Methods for more details on the procedure.

#### KAPACv2.0

We have also implemented a generalized version of our previously published KAPAC tool ^7^. KAPACv2.0, standing for **k**-mer **a**ctivity on **p**olyadenylation site **c**hoice **v**ersion **2.0**, models genome-scale changes in 3’ end usage to infer sequence motifs that can explain differences between samples. In contrast to its first version, KAPACv2.0 can also use both binding sites predicted with position-dependent weight matrices (PWMs) as well as k-mer counts. Also, while the first version of KAPAC was designed to run on sample contrasts, such as knockdown versus control, KAPACv2.0 does not require such contrasts but can be applied to a number of different tissues or a time series. First, we define the relative usage of poly(A) site *p* in sample *s* as *u_p,s_*. KAPAC models the relative usage of poly(A) sites with respect to the mean of all samples as a linear function of the occurrence of PWM binding sites or k-mer counts and the unknown ‘activity’ of these PWMs / k-mers: *log_2_(u_p,s_)* = *N_p,k_* * *A_k,s_* + *c_p_* + *c_s,e_* + ε, where *c_s,e_* is the mean relative usage of the poly(A) site from exon *e* in sample s, *c_p_* is the mean *log_2_* relative usage of poly(A) site *p* across all samples, *N* is the number of binding sites (predicted with the PWM or by k-mer counting) around poly(A) site *p* and ε is the residual error. Finally, *A_k,s_* is the activity of the PWM / k-mer *k* in sample *s*, which determines how much the PWM / k-mer contributes to the relative usage of the poly(A) site. A detailed inference of the model can be found in ref. ^7^. KAPACv2 calculates for every PWMs or k-mer, respectively, a z-score z = *A_k,s_* / *σ_k,s_*, whereas σ are the fitting errors of the activities *A_k,s_*. Background correction and ranking of PWMs / k-mers is done as described for the MAEI approach described above.

### Curation of PWMs of RBPs binding motifs

ATtRACT is a publicly available database of RNA-binding proteins and associated motifs ^44^. On 20 August 2021 we downloaded the set of all available RBP motifs in the format of position-dependent weight matrices (giving the probability of observing any of the four bases at each position of the binding site) as well as their corresponding metadata. We first selected motifs annotated to wild-type *Homo sapiens* proteins obtained with SELEX method. We manually removed duplicate records as well as identical motifs associated with distinct RBPs, as we do not consider them as reliably describing the specificity of individual RBPs. We further trimmed non-informative positions from both ends of each PWM, leaving only the ‘core’ motif with a non-zero information content per position. We did not include PWMs whose core was shorter than 4 and longer than 7 nucleotides. Finally, for each PWM we calculated the total motif entropy and discarded those that were too degenerate (with an entropy higher than 10). This procedure yielded 344 PWMs for the MAPP analyses.

### Density profiles of RBP-specific binding motifs

To gain additional confidence in MAPP’s inferences, we constructed profiles on motif densities around 5’SS, 3’SS and poly(A) sites for exons whose inclusion or 3’ end processing changed most in the respective experiments. We contrasted the motifs of exons whose inclusion level increased, decreased, or was not affected upon change in expression of the corresponding RBP. Specifically, for each plot in Figs. 2 and 3 we selected two groups of the top 200 targets with the highest condition-wise change (in either direction) in alternative splicing as well as alternative polyadenylation based on the average quantified exon inclusion fraction and poly(A) site usage, respectively. Additionally we selected a group of 200 sites with the least change and treated them as a baseline level for motif counts in non-targets. Then, based on the inferred coordinates of binding sites for respective PWMs reported by MotEvo ^45^ in the regions around 3’SS, 5’SS and PAS covered by our sliding windows, we computed the per-position frequency of occurrence of the motif in the proximity of sites from each of the three groups. We then displayed these profiles after mean-based smoothening in the −5/+5nt neighbourhood of each position.

### Selection of ENCODE experiments, reported motifs and k-mers

We have downloaded and analyzed RNA-seq samples linked to 472 knock-down experiments of RBPs, publicly available as a part of the ENCODE project; 16 of these did not pass the quality-control step as defined in the “Datasets” section. We used the remaining 456 data sets for the analysis shown in Fig. 3. Briefly, each ENCODE experiment has been analysed with MAPP in both PWM-based and k-mer-based approaches. For each of the knock-down experiments we selected the ATtRACT PWM associated with the perturbed RBP for which MAPP found the highest statistically significant impact on any of the signals (statistical significance annotated with the “abs” strategy, please see Supplementary Methods) We then noted the ranks of that PWM (among all possible PWMs) in explaining the usage of 3’SS, 5’SS and PAS. The lowest of these three ranks we called the “PWM rank” and this is the key by which the results table is sorted in descending order. For only 14 ENCODE experiments we found that the PWM associated with the perturbed RBP had a rank of maximum 5 (out of 344 curated PWMs). For these, we report the PWM ranks and their impact maps, as inferred by MAPP. We then checked whether the appropriate motif is also recovered in the k-mer mode. For this, we selected the k-mer with the highest statistically significant activity z-score on any of the processing sites (labelled as “1st ranked k-mer”) from each experiment. Alongside with the previously described PWM-based results we report the 1st ranked k-mer and its impact map.

### Data availability

The results generated in this study are available in the supplementary data, which are accessible from Zenodo under doi: https://doi.org/10.5281/zenodo.5789987. The accession numbers for used datasets are available from Supplementary Tables S1 and S2.

### Code availability

The MAPP code is available at https://github.com/gruber-sciencelab/MAPP under the Apache 2.0 open-source license.

