## Supplementary Materials for "Frequent co-regulation of splicing and polyadenylation by RNA-binding proteins inferred with MAPP"

#### 1 Supplementary Figures

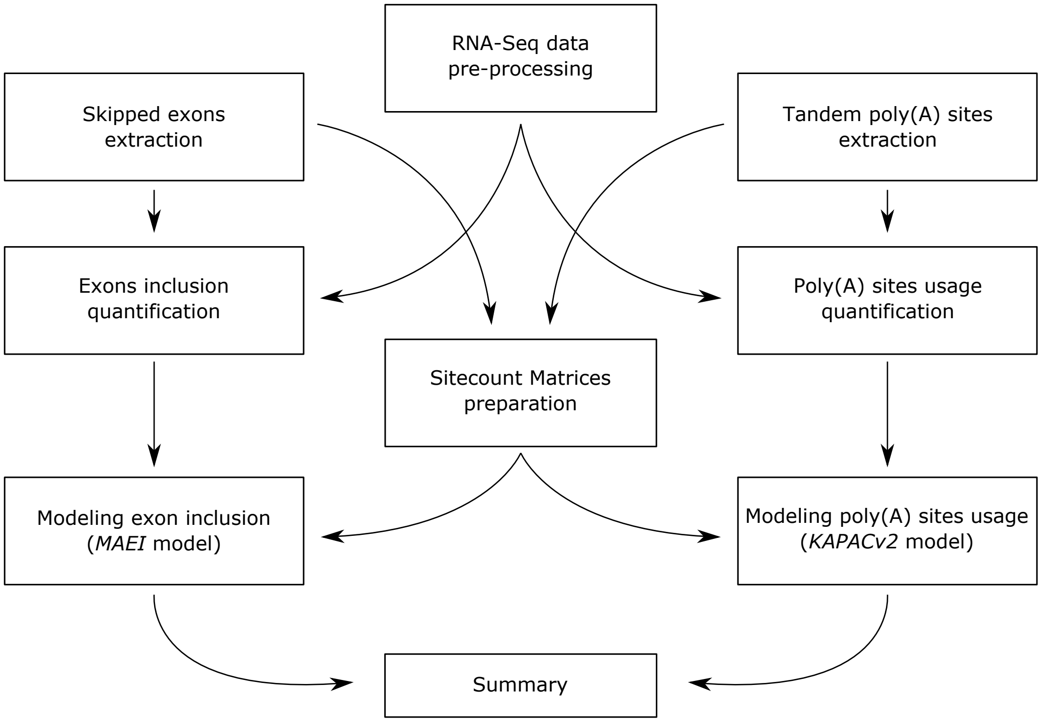


**Figure S1.** High-level overview of MAPP. The pipeline may be decomposed into nine separate functional sub-modules, each of which can be executed individually on its own. Modules on the left-hand side are related to the analysis of alternative splicing whereas right-hand side is dedicated to alternative polyadenylation. Center modules either prepare data for the others or gather and post-process the results. Thus, while running MAPP the execution starts from three independent entry points.

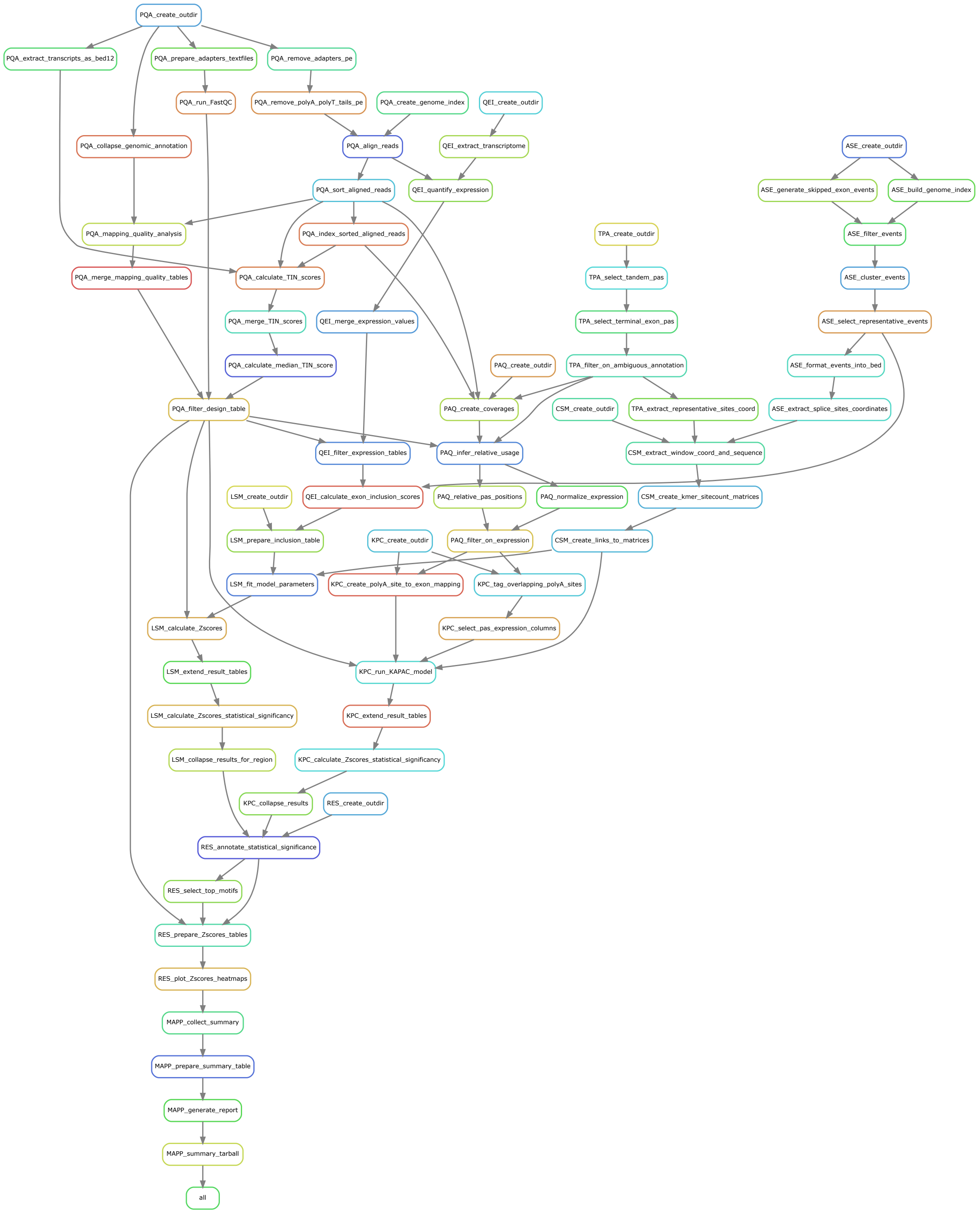


**Figure S2.** Automatically generated snakemake rule graph of the entire MAPP pipeline. Rules of distinct sub-modules of the workflow are prefixed with a three-letter code. Four additional top-level rules are added (prefix: *MAPP*) which generate a compressed HTML-formatted report of the whole analysis run.

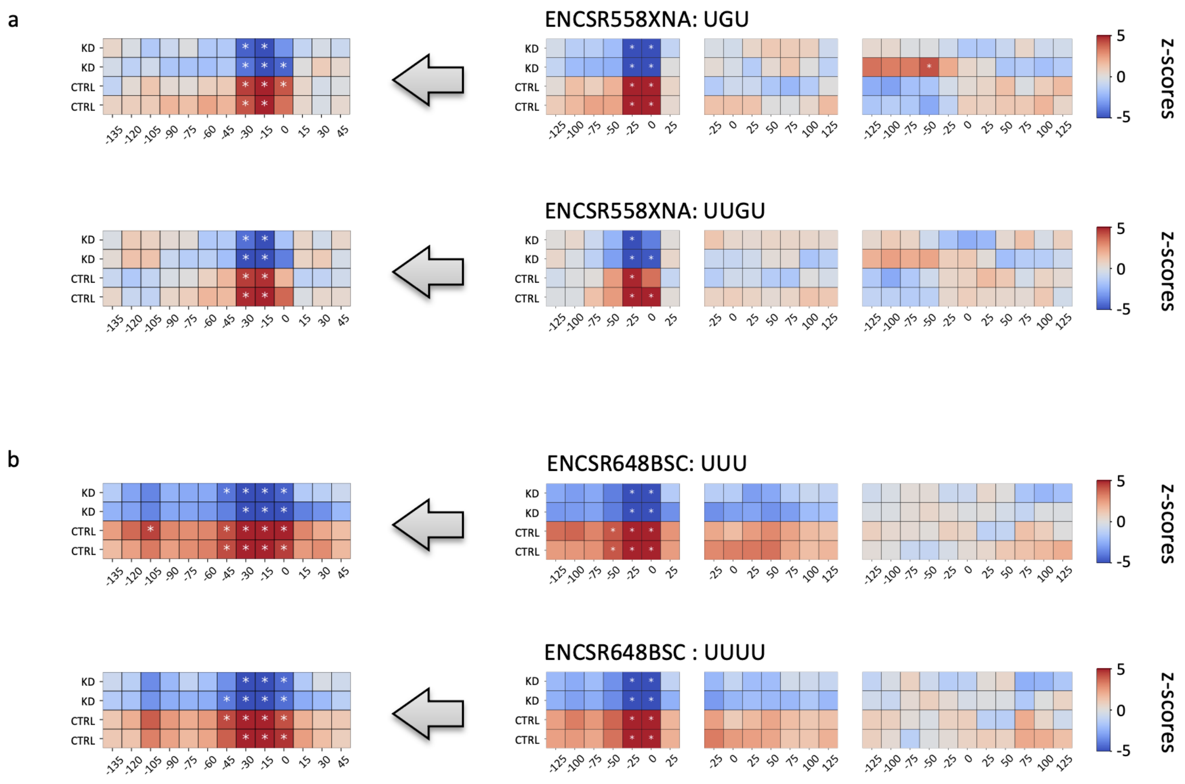


**Figure S3.** Impact maps of top two most significant k-mers reported by MAPP after analyzing two PUF60 knock-down experiments from the ENCODE project. (a) PUF60 knock-down in K5643 cell line. (b) PUF60 knock-down in HepG2 cell line. In order to obtain more fine-grained insight into the position-specific activity of the RBP we rerun our analyses with more narrow sliding windows around 3'SS (30nt in length, slided by 15nt).

### 2 Supplementary Methods

#### 2.1 Data preprocessing

Processing of raw sequencing reads in FASTQ/FASTA format starts with two consecutive runs of cutadapt tool [1]. First, the sequences of adapters are removed and subsequently we trim poly(A) tails. Following that the reads are mapped to both genome and transcriptome using STAR aligner [2], which also - based on the provided resources - builds a genomic index prior aligning the reads. Obtained alignments in BAM format were then sorted and indexed with samtools [3]. If appropriate parameters are set MAPP will also carry out quality control of the data and, prior to the downstream analysis, filter-out such RNA-Seq samples which do not meet specified criteria. For the quality analysis we use metrics provided by RNA-SeQC [4] and TIN score calculated as in RSeQC package [5]. Additionally all samples are handled over to FastQC [6] which generates additional per-sample summary report.
Snakemake rules which belong to this module are marked with a three-letter namespace: *PQA*.

#### 2.2 Selection of cassette exons

We select a set of cassette (also known as 'skipped') exons based solely on the standard ENSEMBL genomic annotation (version: hg38). We first run SUPPA2 [7] to generate all skipped exon events and then filter them for minimal length sufficient for the downstream analyses. We focus only on records annotated as *protein_coding*. Obtained events are further clustered according to a mutual coverage dissimilarity measure:

$$d(e_1, e_2) = 1 - min(\frac{len(e_1 \cap e_2)}{len(e_1)}, \frac{len(e_1 \cap e_2)}{len(e_2)}) \quad (1)$$

Where $e_1$ and $e_2$ denote two exons and with $\cap$ we take only the overlap between then. We applied a hierarhical clustering with a maximum linkage strategy and set 0.05 as a cluster linkage cutoff parameter. From every cluster we selected one event with the highest number of distinct transcripts that support it. Such an event we call a representative exon. For all representative exons we keep the information which transcripts of a given gene include it as well as the list of all transcripts for a given gene (both provided by SUPPA2). We also save the coordinates of 3' and 5' splice-sites of these representatives.
Snakemake rules which belong to this module are marked with a three-letter namespace: *ASE*.

#### 2.3 Extraction of tandem poly(A) sites

We select a set of poly(A) sites which may be classified as proximal/distal within a given terminal exon based on a provided BED-formatted poly(A) site atlas as well as GTF-formatted genomic annotation. Throughout the following study we use PolyASite 2.0 [8] and ENSEMBL annotation, version hg38. We focus on all sites supported by at least one protocol (atlas-specific information) but filter for such which are located only on protein-coding transcripts. We also discard all sites which could be ambiguously annotated to distinct genes. We export coordinates of the resulting tandem polyA sites into a BED-formatted list.
Snakemake rules which belong to this module are marked with a three-letter namespace: *TPA*.

#### 2.4 Quantification of exon inclusion

In order to quantify transcripts' expression based on transcriptomic alignments we run Salmon [9]. Having obtained per-sample, per-transcript TPM-normalized expression values we use the previous information to collapse the scores into a more exon-centric level: for each representative exon (previous section) in every sample we calculate two values: summed up total expression of all transcripts which include this exon ($i_{e,s}$) as well as summed up total expression of all transcripts of a given gene ($t_{e,s}$).
Snakemake rules which belong to this module are marked with a three-letter namespace: *QEI*.

#### 2.5 Quantification of poly(A) sites' expression

In order to quantify the expression of distinct tandem poly(A) sites we employed our previously developed tool: PAQR [10]. The method takes as input genomic alignments of RNA-Seq reads in BAM format and a BED-formatted list of poly(A) sites of interest. It infers expression of distinct sites directly from the coverage profiles in their proximity (please see original publication for more details). As an output it provides TPM-normalized expression of poly(A) sites in all samples and, additionally, a list of their relative position within respective terminal exons. Furthermore, we filter the output table to keep tandem sites only located on such exons for which all their sites are considered as expressed in all analyzed samples.
Snakemake rules which belong to this module are marked with a three-letter namespace: *PAQ*.

#### 2.6 Sitecount Matrices

Both statistical models for discovering regulators of casette exon inclusion and poly(A) sites' usage require tables containing quantified information about binding sites of distinct RNA-binding proteins within a certain distance relative to the site of interest. We call such tables 'sitecount matrices'. In case of alternative splicing analysis we define a range around 3'SS and 5'SS. For the alternative polyadenylation analysis we focus around tandem poly(A) sites. We employ a 'sliding-window' strategy in order to gain a better resolution into the positional-dependent effect of RBPs binding sites on the mRNA maturation process. Thus, the pipeline generates multiple sitecount matrices for 3'SS, 5'SS and poly(A) sites - each one corresponding to a unique window defined relatively to the site of interest. MAPP pipeline can be run in two modes: *kmer-* and *pwm-based*, depending on how the user chooses to generate sitecount matrices. In both cases the module reads in BED-formatted files with the positions of 3'SS and 5'SS of cassette exons as well as locations of tandem poly(A) sites (all previously generated) and produces files with coordinates of distinct windows around them. We extract genomic sequences of the regions encoded in these windows with Pybedtools [11]. In the former mode we sum up the occurances of distinct kmers over the sequence of the whole window (making sure not to overcount short overlapping homomeric subsequences) and the resulting sitecount matrix contains raw counts. In the *pwm-based* approach we utilise MotEvo [12], a probabilistic method which quantifies binding probabilities between nucleotide sequences and distinct motifs (in PWM format). In this case the output sitecount matrices contain posterior probabilities for an RBPs binding event. Regardless of the approach selected the resulting information is stored in a per-exon, per-motif matrix ($N_{e,m}$), for each window separately. Throughout the following study whenever we run MAPP in *kmer-based* mode we count all 3-mers, 4-mers and 5-mers. In case of *pwm-based* analyses we infer binding probabilities of pre-filtered subset of motifs deposited in the ATtRACT databse [13].
Snakemake rules which belong to this module are marked with a three-letter namespace: *CSM*.

#### 2.7 Modeling exons' inclusion (MAEI)

Our aim is to model previously quantified inclusion of cassette exons with the the information stored in sitecount matrices (either raw counts of kmers or binding probabilities for distinct PWMs). Please note that the following procedure is applied to each of the sliding windows separately.

For every exon $e$ we model its inclusion fraction in sample $s$ with a logistic function $\Theta_{e,s}$:

$$f_{e,s} = \frac{i_{e,s}}{t_{e,s}} \sim \Theta_{e,s} = \frac{e^{b_s + c_e + N_{e,m} \times A_{m,s}}}{1 + e^{b_s + c_e + N_{e,m} \times A_{m,s}}} \tag{2}$$

where $b_s$ and $c_e$ are parameters of the model which account for the baseline inclusion level of all exons in a given sample $s$ and the baseline inclusion level of $e$ across all samples, respectively. We call $A_{m,s}$ the activity of a given motif $m$ in $s$ and this is the parameter which accounts for the effect of short sequence motifs on the differential exon inclusion process.

We calculate the probability of observing all the quantified data obtained from the RNA-Seq experiment as a product of consecutive Bernoulli trials where exon inclusion is treated as "success" and exclusion as "failure". Under that model we write down a likelihood function:

$$P(D|M) = \prod_{s,e} \Theta_{e,s}^{i_{e,s}} \times (1 - \Theta_{e,s})^{t_{e,s} - i_{e,s}} \tag{3}$$

In order to mitigate the dominant effect of highly expressed genes on the likelihood we introduced a scaling prefactor:

$$R = \frac{C}{X} \times \frac{\langle t_e \rangle}{\langle t_e \rangle + t_{crit}} \tag{4}$$

where $C = \sum_{e,s} t_{e,s}$ and $X = \sum_{e,s} \frac{\langle t_e \rangle}{\langle t_e \rangle + t_{crit}}$, the averages are taken over all samples $s$. Introducing the $t_{crit}$ hyperparameter allows us to bound to weights the likelihood contribution from distinct exons. Exons originating from highly expressed genes saturate the prefactor thus the whole likelihood is not overtaken by gene expression level. We set that $t_{crit}$ is always inferred from the data as the median of the distribution of $\langle t_e \rangle$ (averaged across all samples).

Finally we write the expression for log-likelihood of our model as:

$$LL = \sum_{s,e} R \times [f_{e,s} \times (b_s + c_e + N_{e,m} \times A_{m,s}) - \log(1 + e^{b_s + c_e + N_{e,m} \times A_{m,s}})] \tag{5}$$

Our aim is to find maximum likelihood estimates of the model. We optimize it by an EM algorithm where we iteratively calculate partial derivatives with respect to model parameters ($\frac{\partial LL}{\partial A_s}, \frac{\partial LL}{\partial b_s}, \frac{\partial LL}{\partial c_e}$), update their values and re-calculate the likelihood until it converges.

We obtain standard deviations of motif activities ($A_{m,s}$) from the Hessian matrix of the log-likelihood function at its optimum. Its negative inverse is an estimator of the covariance matrix of the model parameters. We use these estimates to standardize the activities: for every sample $s$ we calculate a per-sample motif activity z-score: $Z_{m,s} = \frac{A_{m,s}}{\sigma_{A_{m,s}}}$.

Motif activity z-scores are then background corrected in a second statistical model. In order to rigurously detect which motifs may be considered as outliers we fit a mixture of a uniform and normal components to the distribution of per-sample z-scores:

$$P(D|M) = \rho \times \frac{1}{\max Z_{m,s} - \min Z_{m,s}} + (1 - \rho) \times \frac{1}{\sqrt{2\pi}\sigma} \times e^{\frac{-(Z_{m,s} - \mu)^2}{2\sigma^2}} \tag{6}$$

Max and min functions are applies over all motifs $m$ for a given sample $s$.

Similarly as previously, we find the maximum likelihood estimates for the parameters of the model ($\rho, \mu, \sigma$) by converging the log-likelihood with an EM algorithm. We later standardize per-sample motif activity z-scores by the parameters of the inferred Gaussian background distribution: $Z^{\#}_{m,s} = \frac{Z_{m,s} - \mu}{\sigma}$. Such scores are later transformed into p-values of a standard normal distribution. In order to assess statistical significance at $\alpha = 0.05$ level we also apply a Bonferonni correction where we adjust by the total number of motifs. Thus for every motif $m$ in every sample $s$ (for every sliding window separately) we get $p_{m,s}$.

Snakemake rules which belong to this module are marked with a three-letter namespace: *LSM*.

#### 2.8 Modeling poly(A) site usage (KAPACv2.0)

For the needs of this project we have developed an advanced version, KAPACv2.0, of our previously published KAPAC approach [10]. KAPACv2.0 is capable of running based on both, binding sites predicted from position weight matrices as well as based on k-mer counts. Also, it can be applied to any type of samples, such as a number of different tissues or a time series and is in contrast to KAPAC not limited to sample contrasts, such as knock-down versus control samples. The KAPACv2.0 approach is summarized in the Methods section of the main article.

Snakemake rules which belong to this module are marked with a three-letter namespace: *KPC*.

#### 2.9 Analysis summary

Following both statistical models and having inferred motif activites ($A_{m,s}$), their z-scores ($Z_{m,s}$) and p-values ($p_{m,s}$) in all samples and in every window we proceed to summarize the analysis, select those results which we consider statistically significant and visualise them with heatmaps of activity z-scores, which we refer to as 'Impact Maps'. In the last module of the workflow we implemented two distinct strategies to filter motifs based on statistical significance, applicable to different analyses types. In the *avg* mode we call a given motif $m$ statistically significant if and only if there exist a window $w$ within which for at least half of the samples $p^w_{m,s}$ are below a previously defined cutoff (0.05). This strategy is designed for common comparative analyses of two biological conditions, each being sequenced in multiple replicates. The other approach - *max* - requires $m$ to be called as statistically significant in only one sample in order to annotate significance to the whole motif. The rationale behind this was to provide an insightful way of investigating datasets which consist of multiple distinct conditions. For every motif called as statistically significant we plot an Impact Map as a visual summary of that motifs activity on exon inclusion and poly(A) site usage.
Snakemake rules which belong to this module are marked with a three-letter namespace: *RES*.

#### 2.10 Final report

At the end of the workflow we designed a few additional steps which prepare an output directory with the most important text files and tables as well as a summary report in HTML format. Summary directory is also compressed into .tar.gz format to facilitate reproducibility and usability: users may exchange their results as well as run configurations easily. The final HTML report contains a sorted table of motifs whose activity z-scores were reported to be statistically significant in at least one sliding window around any of the processing sites. The columns of that table include: motif ID, sequence logo (in case of a PWM-based MAPP execution), three maximum activity z-scores (with the maximum taken over all sliding windows around each of the analyzed sites: 3'SS, 5'SS, PAS), ranking score (set to an average of the three aforementioned z-scores) and the Impact Map.

Top-level snakemake rules related to the final summary are marked with a four-letter namespace: *MAPP*.
